# RNA N6-adenine methylation dynamics impact *Hyaloperonospora arabidopsidis* resistance in *Arabidopsis*

**DOI:** 10.1101/2024.01.30.577950

**Authors:** Leonardo Furci, Jérémy Berthelier, Hidetoshi Saze

**Author notes:** Authors contributed equally (Leonardo Furci, Jérémy Berthelier).

## Abstract

In plants, epitranscriptomic mark N6-adenine methylation (m6A) is dynamically regulated by (a)biotic stressors. However, there is limited knowledge on m6A dynamics at single nucleotide resolution on specific RNA molecule under stresses, and their role in environmental adaptation. By using Oxford Nanopore Technology direct RNA sequencing (ONT-DRS) and a neural network model, here we show transcript-specific dynamics of m6A modification at single nucleotide resolution by biotic stress during *Hyaloperonospera arabidopsidis* (*Hpa*) infection in *Arabidopsis*. In wild-type seedlings, pathogen infection causes significant reduction of global m6A ratios, which correlates with activation of m6A-modified transcripts. Defect of m6A deposition in the m6A mutant *hakai-1* mimics m6A reduction from *Hpa* infection at ∼70% of sites, resulting in constitutive overexpression of basal defence genes and enhanced resistance against the pathogen. Our results demonstrate that m6A dynamics impact defence response against pathogen, providing a promising target for future crop improvement strategies.

## Introduction

Epitranscriptome comprises over 170 RNA modifications, with N6-adenine methylation (m6A) being the most extensively studied and abundant in eukaryotes, particularly in mRNA (Shinde et al., 2023). In the model plant *Arabidopsis thaliana*, m6A is deposited by a complex of methyltransferases mRNA adenosine methylase A (MTA, homolog of mammalian METTL3) and methyltransferase B (MTB, homolog of mammalian METTL14), along with cofactors VIRILIZER (VIR, homolog of mammalian VIRMA), E3 ubiquitin ligase AtHAKAI (homolog of mammalian HAKAI) and FKBP12 INTERACTING PROTEIN 37 (FIP37, homolog of mammalian WTAP) (Růžička et al., 2017; Shen, 2023). In mRNA, m6A is deposited at the RRACH (R = A/G, H = A/U/C) consensus motif, and is mostly prevalent near start/stop codons and at 3’ untranslated regions (3′-UTRs) of mature mRNAs (Shinde *et al*., 2023). Several studies in different eukaryotic systems have shown that m6A modulates mRNAs fate in cells, by affecting mRNA stability and half-life, alternative splicing, mRNA translation, nucleo-cytoplasmic export rates, and m6A-mediated phase separation (Zaccara et al., 2019). Removal of m6A is mediated by α-ketoglutarate-dependent dioxygenase homolog (ALKBH) proteins, such as ALKBH9B and ALKBH10B (Shinde *et al*., 2023) in *Arabidopsis* (Mielecki et al., 2012). Recent studies reveal that abiotic environmental cues such as salt, cold or heat stresses dynamically modulate N6-adenine (de)methylation (Hu et al., 2021; Wang et al., 2022a). In wild-type (Wt) plants, heat stress application increased m6A levels, while cold treatment reduced them, particularly at 3′-UTR of mRNAs (Wang *et al*., 2022a; Wang et al., 2023). Additionally, loss-of-function mutants for the core components of m6A deposition displayed salt stress (Hu *et al*., 2021) and cold stress (Wang *et al*., 2023) hypersensitivity, whereas m6A demethylase mutant *alkbh10b-1* was impaired in thermotolerance and flower development (Duan et al., 2017; Wang *et al*., 2022a), underlining the significance of m6A dynamics in plant-environment interactions. More recently, a role of m6A in the context of pathogens resistance in *Arabidopsis* has been suggested (Prall et al., 2023). However, quantitative analyses of m6A dynamics at single-nucleotide resolution on single mRNA molecules during biotic stress and plant defence responses remain unresolved due to technical limitations.

## Results

To investigate the role of m6A in plant basal defence to pathogens, we inoculated Wt, *hakai-1*, *hakai-2* and *mta* (Růžička *et al*., 2017) *Arabidopsis* seedlings (Col-0 accession) with biotrophic oomycete *Hyaloperonospera arabidopsidis* WAC09 (*Hpa*), biotrophic bacteria *Pseudomonas syringae* pv *tomato* DC3000 (*Pst*) and necrotrophic fungus *Plectosphaerella cucumerina* PMM (*Pc*), and assessed their disease resistance. Compared to Wt, mutants *hakai-1* and *hakai-2* exhibited a significant reduction in *Hpa* leaf colonization (Chi-squared tests, *p* <0.05), indicating an elevated resistance, while the *mta* mutant showed a small reduction in *Hpa* colonization that was not significantly different (Figure 1A). All three mutants showed a mild reduction in *Pst* colonization, albeit not significant, and displayed no differences in resistance/susceptibility against *Pc* (Figure 1A). To dissect transcriptional responses of Wt and m6A deposition mutants to *Hpa* inoculation, we performed transcriptomic analysis of Mock- and *Hpa*-treated Wt, *hakai-1*, *hakai-2* and *mta* seedlings at 48 hours (48h) after treatment using short-read Illumina RNA-seq. Differential gene expression (DGE) analysis of *Hpa*-treated Wt seedlings identified 947 upregulated and 533 downregulated genes compared to Mock-treated control. Upregulated genes showed significant enrichment of gene ontology (GO) biological processes (BP) terms related to both early-stage defence responses to *Hpa,* such as reactive oxygen species production or callose deposition, and late-stage ones, such as salicylic acid-dependent responses (Supplemental Figure 1), as previously reported for this time-frame following pathogen inoculation (Furci et al., 2019). A comparison between Wt and mutants revealed a subset of 188 differentially expressed genes (DEGs) specifically upregulated in *Hpa*-resistant *hakai-1* and *hakai-2* mutants under basal (Mock) conditions, significantly enriched for GO terms for BP related to biotic stress responses (Figure 1B and Supplemental File 1). On the other hand, downregulated DEGs in any condition showed no GO terms enrichments related to basal defences (Figure 1B and Supplemental File 1). Focusing on *Hpa*-inducible genes of Wt that showed differential induction in *hakai* mutants, we identified three major clusters based on their expression patterns: *Hpa*-inducible genes showing constitutive upregulation in *hakai* mutants (Cluster I); *Hpa*-inducible genes showing enhanced induction in *hakai* mutants (Cluster II, similar to the transcriptional responsiveness of primed plants); *Hpa*-inducible genes showing a combination of constitutive overexpression and enhanced induction (Cluster III) (Figure 1C). Notably, only genes in Cluster I and III showed enrichment for defence-related GO terms (Figure 1C and Supplemental File 1), particularly related to salicylic acid-dependent defence pathway required for *Hpa* resistance (Herlihy et al., 2019). Together, these results indicate that increased basal resistance to *Hpa* in *hakai* mutants correlates with the constitutive overexpression of defence-related genes.

**Figure 1:**
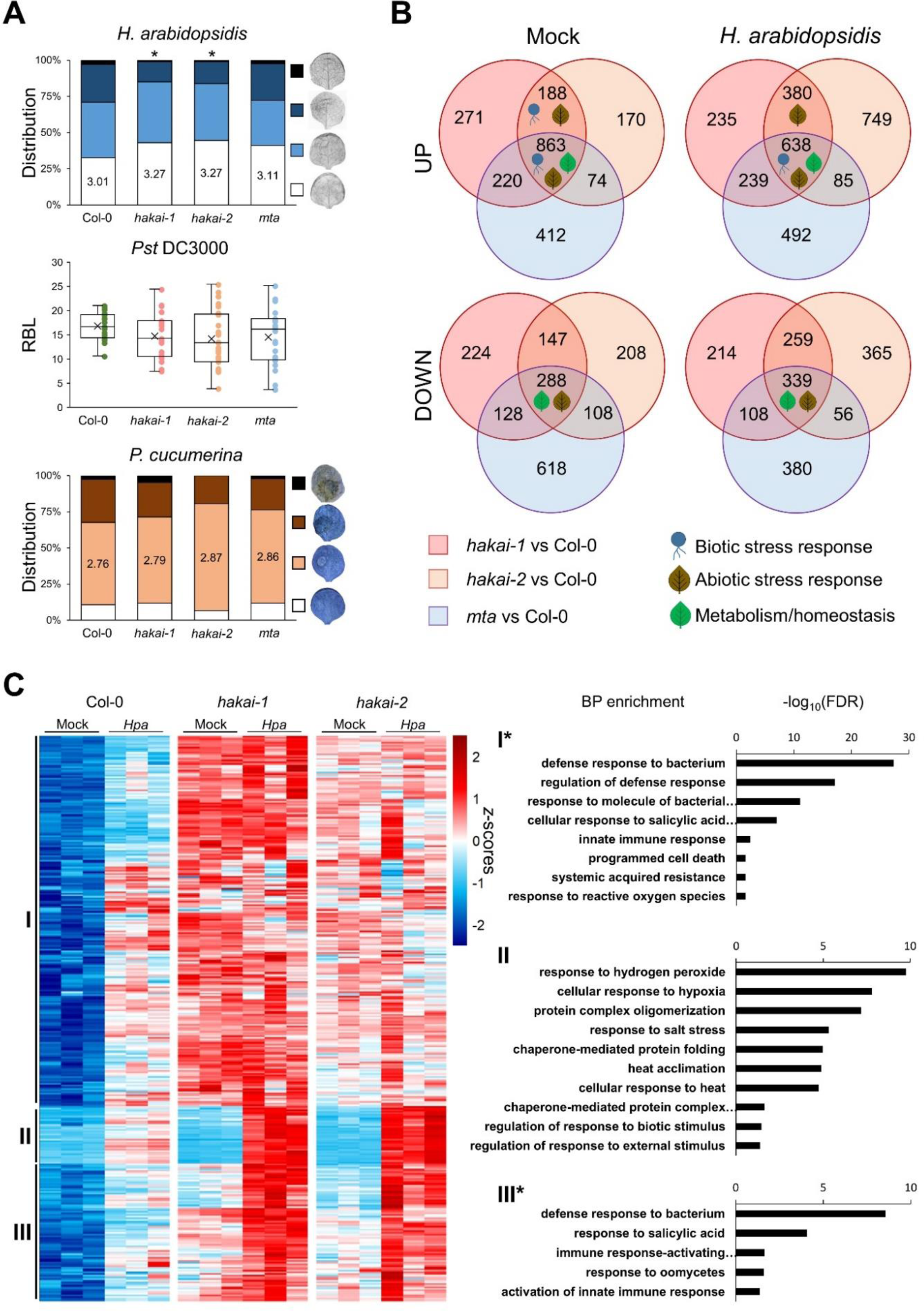
Phenotypical and transcriptional characterization of m6A deposition mutants. **A:** Basal resistance phenotype of *hakai-1*, *hakai-2* and *mta* mutants against different pathogens. Top graph shows distribution of infection classes for *Hpa-*treated leaves from each genotype, ranging from least infected (white), to most infected (black, examples shown on the right). Statistically significant differences in *Hpa* leaf colonization between Wt and mutants were assessed by Chi-squared test (n =150+). Middle graph shows quantification of relative bioluminescence (RBL) from transgenic *Pst* colonization (x indicates mean RBL for each genotype). Statistically significant differences in *Pst* colonization between each mutant and Wt were assessed by 2-tailed Student’s T-test (n =20+). Bottom graph shows distribution of severity of necrotic lesions for *Pc* drop-inoculated leaves, ranging from no lesion (white) to completely necrotized leaves (black, examples shown on the right). Statistically significant differences in *Pc* necrotic lesion category distribution between Wt and mutants were assessed by Chi-squared test (n =35+). Asterisks indicate *p* < 0.05. **B:** Distribution and overlaps of up- or down-regulated DEGs in m6A deposition mutants. Each circle represents DEGs in pairwise comparisons between the indicated mutant and Wt at the appropriate condition (Mock of *Hpa*). Comprehensive list of enriched BP terms is provided in Supplemental File 1. **C:** Hierarchical clustering (Ward’s method) of *Hpa*-inducible genes in Wt seedlings showing differential induction in both *hakai* mutants. For each cluster (I, II, III), GO terms enrichment for BP are shown on the right. Asterisks indicate clusters that contain additional GO terms not related to basal defences (full list provided in Supplemental File 1).

To investigate the dynamics of m6A and its impact on transcription during *Hpa* infection, we sequenced mRNA from Mock and *Hpa*-treated Wt and *hakai-1* seedlings using ONT-DRS. We exploited the DENA neural network tool that demonstrated identification of 90% of miCLIP-detected m6A sites and quantification of m6A ratio at single-nucleotide resolution in the RRACH context from ONT-DRS raw signals (Qin et al., 2022). Analysis of the global distribution of m6A at RRACH sites across all conditions revealed a significant reduction in average m6A during *Hpa* infection in Wt seedling compared to Mock-treated ones (Figure 2A). Similarly, *hakai-1* mutation led to a significant reduction in m6A averages in both Mock- and *Hpa*-treated mutant seedlings (Figure 2A). These results align with a previous report that observed a partial reduction of m6A in *hakai* mutants using two-dimensional thin layer chromatography, in comparison to other mutants of m6A deposition like *vir* or *fip37* (Růžička *et al*., 2017). The number of transcript-specific RRACH sites in each sample meeting the minimum coverage threshold (50 reads) varied proportionally to each sample’s read depth, as expected (Figure 2A, Supplemental Figure 2A). Hence, we focused on a subset of 8691 m6A sites shared across all four conditions to ensure a more robust quantification of m6A changes by pathogen treatment or mutation (Supplemental Figure 2B). Across transcript regions, we confirmed an overall higher m6A levels around 3′-UTRs (Figure 2B), as previously reported across different model species (Shinde *et al*., 2023; Wang et al., 2022b). Notably, we found that *Hpa* treatment of Wt resulted in a significant reduction in m6A, particularly at CDS region of transcripts (Figure 2B). In contrast, *hakai-1* mRNAs showed significant reductions in m6A across entire transcripts, particularly at 3′-UTRs compared to Wt, without strong impact of *Hpa* treatment on m6A levels (Figure 2B). These results demonstrate the presence of dynamic m6A regulation during the defence response.

**Figure 2:**
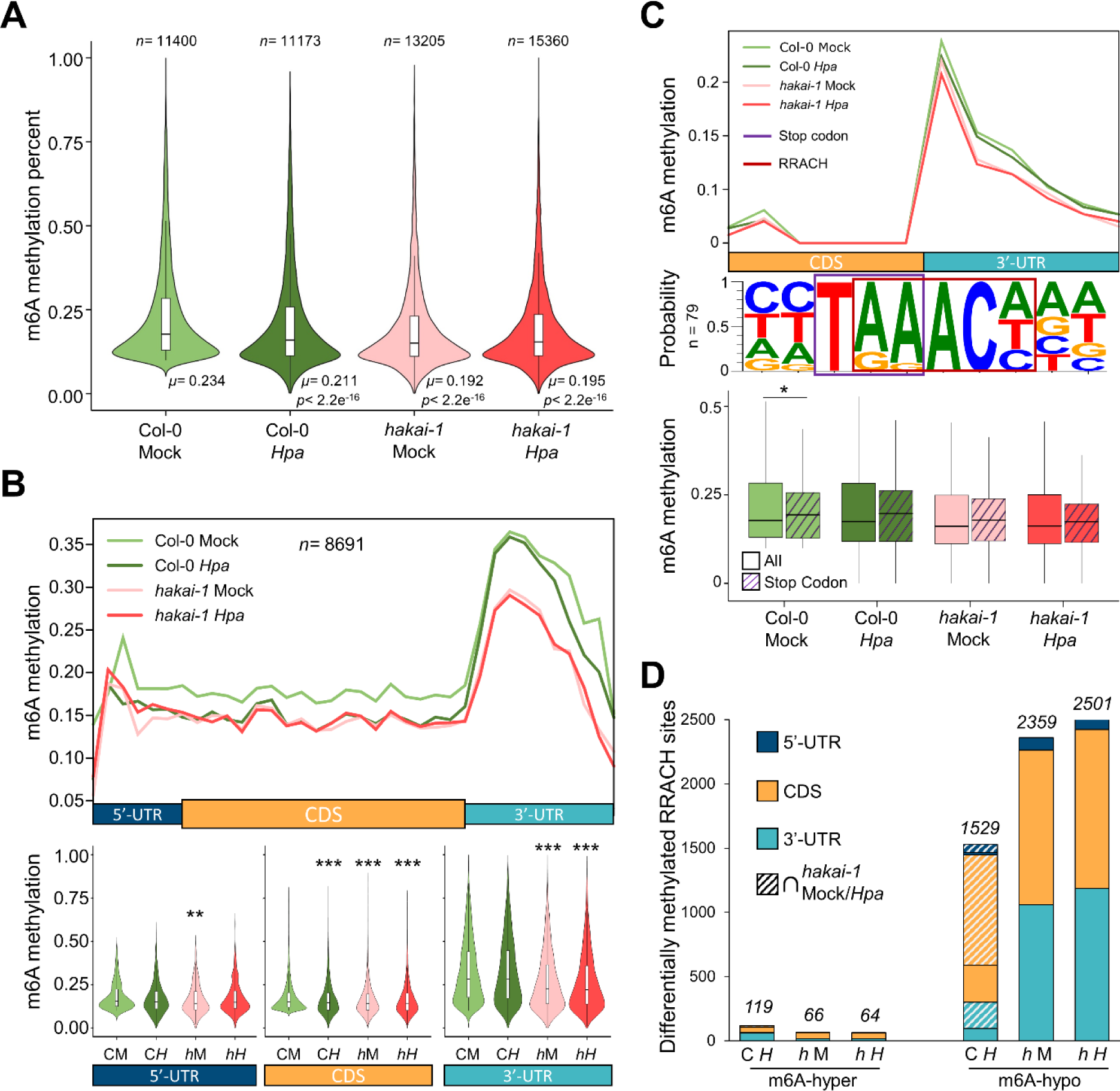
Dynamics of m6A in Wt and *hakai-1* Arabidopsis seedlings upon *Hpa* infection. **A:** Global distribution and average m6A percentages in Mock- and *Hpa*-treated Arabidopsis seedlings. Statistically significant differences in distribution were assessed by Student’s T-test in pairwise comparisons with Mock-treated Wt (*μ* = mean; n. s. = not significant) **B:** Top panel: Mean m6A methylation ratio (50bp resolution) over transcript regions for shared m6A sites (*n*= 8691, Supplemental Figure 2) across conditions. Bottom panel: distribution of methylation ratios for shared m6A sites in each transcript region. Statistically significant differences in distribution were assessed by Student’s T-test in pairwise comparisons with Mock-treated Wt. **, *p* < 0.01; ***, *p* < 0.001. **C:** Top panel: mean m6A methylation ratio at 1bp resolution around transcripts stop-codon. Mid panel: motif enrichment of stop-codon terminal m6A sites. Purple box indicates stop-codon and red box indicates RRACH motif. Bottom panel: m6A methylation ratios for stop-codon terminal m6As (purple stripes, *n*= 79) compared to all m6A (no stripes, *n*= 8691). Statistically significant differences in distribution were assessed by Student’s T-test. *, *p* < 0.05. **D:** Number of differentially methylated m6A sites compared to *Hpa*-treated Wt (Fisher’s exact test), separated by transcript region location. Striped bars indicate differentially methylated m6A sites shared between *Hpa*-treated Wt and Mock-/*Hpa*-treated *hakai-1* mutant. C= Col-0; *h* = *hakai-1*; M = Mock treatment; *H* = *Hpa* treatment.

While previous studies identified an enriched peak of m6A downstream of CDS using immunoprecipitation techniques, they could not resolve the exact m6A positions at single nucleotide resolution (Dominissini et al., 2012; Prall *et al*., 2023). By separating m6A profiles across each of the 12 unique RRACH motif combinations, we found overall higher m6A ratios at RRACH motifs starting with AA or GA (Supplemental Figure 3A). Intriguingly, visualizing the distribution of m6A at 1bp resolution revealed the highest m6A peak specifically at one nucleotide downstream of CDS stop-codon (Supplemental Figure 3B). For these stop-codon terminal m6As, the last two AA/GA/AG nucleotides of the stop-codon triplet have been co-opted as first two elements of the m6A RRACH motif (Figure 2C, middle panel, Supplemental Figure 3C). Additionally, we found that in *Arabidopsis* stop-codon terminal m6As showed a preference for TAA stop-codon trinucleotide (Supplemental Figure 3C), in contrast to the TGA trinucleotide reported in human or mouse cells (Luo et al., 2022). Transcripts possessing stop-codon terminal m6A had the highest significant GO enrichment for BP related to translation, of which all genes coding for ribosomal proteins (Supplemental Figure 3D). Further investigations are needed to elucidate the functional relevance of the stop-codon terminal m6A in transcription and translation control. Quantification of differentially methylated adenines in *Hpa-*treated Wt seedling compared to Mock revealed 119 significantly hyper-methylated (m6A-hyper) or 1529 hypo-methylated (m6A-hypo) m6A sites (Figure 2D). Consistently, we found *Hpa*-dependent upregulation of HAKAI-interacting zinc finger protein 1 (*HIZ1*, Supplemental Figure 4) in Wt seedlings, which is known to downregulate HAKAI-dependent methylase activity, causing m6A reduction when overexpressed (Zhang et al., 2022). These results demonstrate, for the first time, dynamic modulation of m6A in Wt seedlings following *Hpa* infection, predominantly leading to reduction of m6A levels at specific mRNA transcripts. As expected, more m6A-hypo sites were found in *hakai-1* mutant in both Mock- and *Hpa*-treated samples (2359 and 2501, respectively), with a larger proportion occurring at the 3′-UTR, compared to m6A-hypo caused by *Hpa* infection in Wt plants (Figure 2D, Supplemental File 2). Notably, 69.2% of m6A-hypo sites in Wt following *Hpa* infection were also hypomethylated in either Mock- or *Hpa*-treated *hakai-1* mutant, compared to Mock-treated Wt (Figure 2D). This result indicates that *hakai-1* mutation mimics most of m6A-hypomethylation caused by *Hpa*-derived epitranscriptional reprogramming in Wt.

Two of the best-characterized roles of m6A in post-transcriptional regulation of RNA are the modulation of alternative splicing and mRNA abundance (Shinde *et al*., 2023). To investigate the effects of m6A changes induced by *Hpa* infection or *hakai* mutation in alternative splicing, we conducted differential transcript usage (DTU) analysis using short-read RNA-seq data of Mock- and *Hpa*-treated Wt and *hakai-1* seedlings. We next crossed DTU results with isoform-specific ONT-DRS m6A calling to pinpoint changes in isoform usage that could be linked by m6A modulation. Overall, *Hpa* infection had the largest effect on DTU, with a total of 268 isoforms for 153 genes showing significant change in isoform usage, whereas only 175 and 183 isoforms for 98 and 106 genes were identified in Mock- and *Hpa*-treated *hakai-1* mutant, respectively, compared to Mock-treated Wt (Supplemental Figure 5). For most genes however, no m6A sites were identified, or data was limited to the main isoform (Supplemental Figure 5). This is most likely due to lower transcript abundance of alternative isoforms, making detection of m6A sites above the minimum coverage threshold (50 ONT-DRS reads/site) more difficult. Nonetheless, we could identify a few genes showing both DTU between two or more isoforms and at least a single differentially methylated m6A site at any isoform (Figure 3A). For these genes, the reduction in m6A at different transcript regions generally correlated with an increased isoform expression in each condition (Figure 3A). For the gene *YUELAO 2* (*YL2*, *At5G25280*), the different proportions of overexpression for isoforms 1 and 2 in *Hpa*-treated *hakai-1* (compared to Mock-treated Wt) correlated with m6A changes at region-specific sites detected only in either isoform (5′-UTR for isoform 1 and CDS for isoform 2, Figure 3B). Additionally, for the most-overexpressed isoform 1, a shared m6A site at the 3′-UTR was specifically hypomethylated in *Hpa*-treated *hakai-1* at this isoform (Figure 3B, purple box). We then shifted our focus to the role of m6A dynamics on regulation of mRNA abundance by combining DGE analysis with ONT-DRS differential m6A calling. We identified 41 genes showing both differential transcriptional responsiveness to *Hpa* in the *hakai-1* mutant (Figure 1C) and significant m6A-hypo or m6A-hyper methylation in at least a single adenine of their transcript in any condition (Figure 3C). For most of these genes, transcriptional activation by *Hpa* in Wt corresponded with a dynamic reduction in m6A in their transcripts (Figure 3C). Interestingly, in *hakai-1*, these genes showed lower m6A at basal conditions (Mock) which correlated with constitutive transcriptional upregulation (Figure 3C). In human cells, deposition of m6A increases mRNA decay by YTHDF2-mediated CCR4/NOT degradation (Wang *et al*., 2022b), suggesting a negative correlation between m6A and transcript abundance. For the subset of genes, our results suggest that the *hakai-1* mutant at basal condition can mimic the *Hpa*-induced m6A reduction associated with transcriptional activation in Wt plants. The 41 targets include many genes previously reported to be involved in pathogen response (Supplemental File 3), such as *MITOGEN-ACTIVATED PROTEIN KINASE 3* (*MPK3*, At3G45640), a well-characterized kinase involved in innate immunity downstream signalling (Sun and Zhang, 2022), and *EARLY RESPONSE TO DEHYDRATION 6* (ERD6, At1G08930), a putative sucrose transporter involved in *AZELAIC ACID INDUCED 1*(AZI1)-dependent priming of salicylic acid induction and systemic acquired resistance (Wang et al., 2016) (Figure 3D, Supplemental File 2). For both genes, m6A reduction localized at multiple sites at CDS and 3′-UTR of their transcript and was proportional to increased gene expression levels caused by *Hpa* infection or *hakai-1* mutation (Figure 3D). In yeast, a causal link between reduction of a single m6A site in the 3′-UTR of meiotic regulator *REGULATOR OF MEIOSIS 1* (*scRME1*) and increase in its mRNA abundance has been reported (Bushkin et al., 2019). Overall, our results suggest a regulation of transcriptional activation of defence genes and pathogen response by modulation of transcripts’ m6A levels at specific sites, which might involve inhibition of mRNA decay activity and stabilization of transcripts.

**Figure 3:**
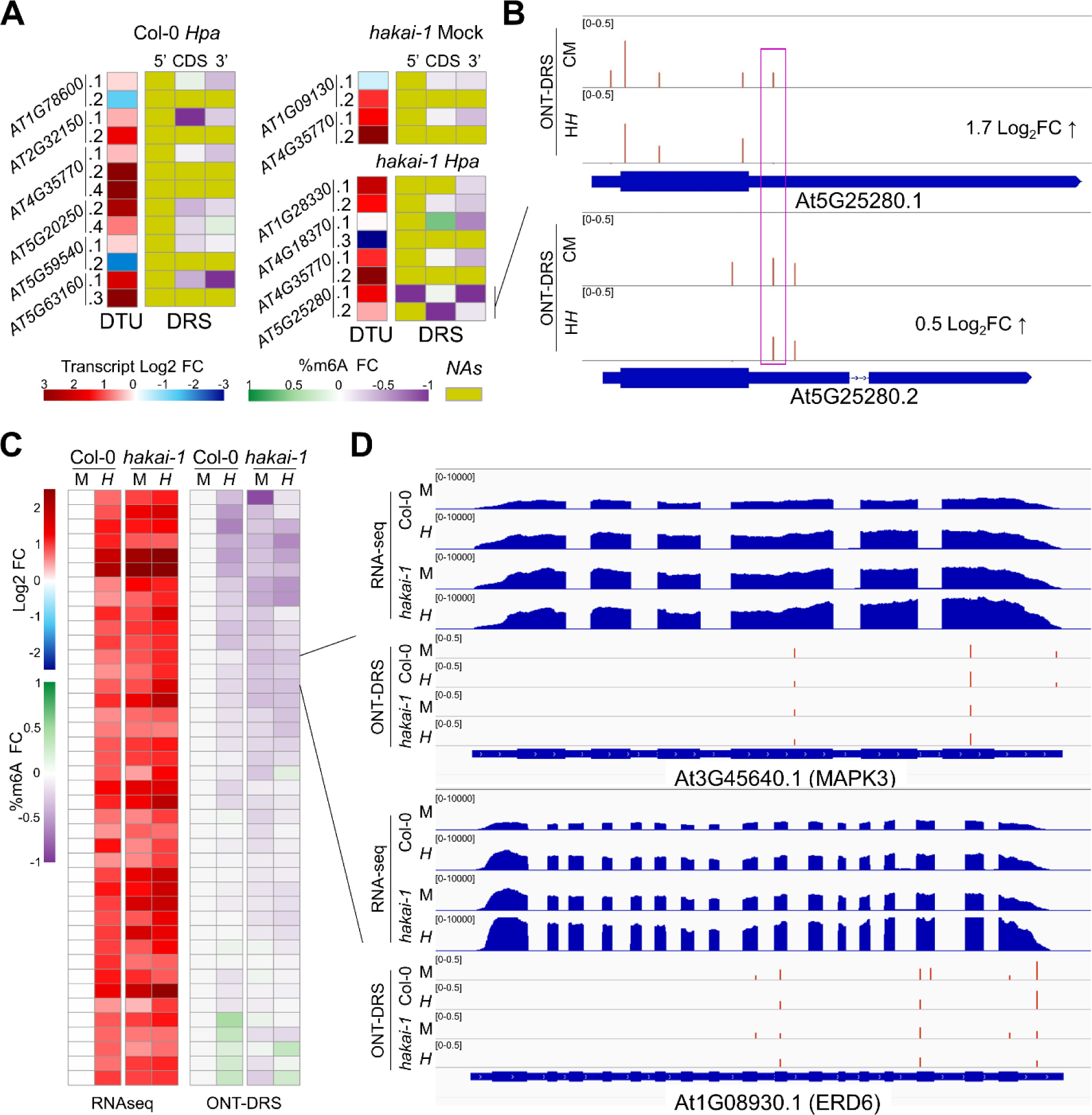
Gene-specific epitranscriptomic regulation of alternative splicing and mRNA accumulation. **A:** Correlation between differential transcript usage (DTU) and m6A changes at selected genes. DTU heatmap displays log_2_ fold changes in isoform expression; DRS heatmap shows fold changes in m6A percentage across transcript region of each isoform (NA = no data available). **B:** Example of transcript-specific m6A changes for differentially spliced At5G25280.1 and At5G25280.2 isoforms in *Hpa*-treated *hakai*-1 vs Mock-treated Wt. ONT-DRS tracks display m6A percentages at identified m6A sites. **C:** Correlation between transcriptional changes and m6A changes of 41 candidate genes. RNA-seq heatmap displays log_2_ fold changes in gene expression; ONT-DRS heatmap shows fold changes in m6A percentage averaged across each transcript. **D:** Browser view of representative candidates *MAPK3* and *ERD6* (with corresponding heatmap cells indicated in Figure 1C). RNA-seq tracks display normalized read coverages; ONT-DRS tracks display m6A percentages at identified m6A sites. C= Col-0; *h* = *hakai-1*; M = Mock treatment; *H* = *Hpa* treatment.

In conclusion, our study provides a first insight into Arabidopsis m6A dynamics in response to biotic stress during pathogen infection. In Wt, epitranscriptional reprogramming following *Hpa* treatment coincides with m6A reduction at specific m6A sites (Figure 2B). In the m6A defective mutant *hakai*, around 70% of *Hpa*-derived m6A-hypo sites in Wt are hypomethylated in both Mock and *Hpa* conditions, suggesting a role of HAKAI in mediating m6A dynamics in response to *Hpa* infection. Increased mRNA accumulation in a subset of *Hpa*-inducible defence-related genes in Wt correlates with transcript-wide m6A reduction (Figure 2C), consistent with previous reports in different model organisms (Bushkin *et al*., 2019; Wang *et al*., 2022b). In the *hakai* mutant, these genes possess lower transcript-wide m6A ratios at basal state, resulting in global constitutive overexpression of biotic stress response genes (Figure 2C and Figure 1B-C) and increased resistance (Figure 1A). Given recent advances in methodologies for targeted m6A mRNA editing (Fang et al., 2023), our findings provide a promising avenue for novel methods of crop protection against pathogens and future crop improvement programs.

## Methods

### Plant materials and growth conditions

The *hakai-1* line (GABI_217A12) was obtained from ABRC (Ohio, US) and genotyped to retrieve homozygous plants. The *hakai-2* and *mta* seeds were kindly provided by Prof. Rupert G. Fray. The *hakai-2* line is homozygous for CRISPR-induced point mutation in *HAKAI,* as described previously (Růžička *et al*., 2017). The mutant *mta* corresponds to homozygous SALK_074069 line complemented with *ABI3:MTA*, to rescue the embryo-lethal phenotype of homozygous T-DNA insertion, as previously described (Bodi et al., 2012). All mutants are in Col-0 background.

Seeds of Wt and mutants were stratified in water for 4 days in darkness at 4°C before sowing in soil. Seedlings were grown for 3 weeks under short-day conditions (8.5 h light/15.5 h dark, 21°C, 60% RH, ∼125μmol s^−1^ m^−1^ light intensity) before pathogen inoculations.

### Pathogen resistance assays

For *Hpa* infection, seedlings were spray-inoculated with a suspension of asexual conidia in sterile water at a density of 10^5^ spores/ml, and kept at 100% RH to promote infection. *Hpa* colonization was quantified six days post-inoculation by microscopic scoring of leaves, as described previously (Berthelier et al., 2023; Furci *et al*., 2019). Trypan blue-stained leaves (Asai et al., 2015; Furci *et al*., 2019) were analyzed with a stereomicroscope (KL200 LED, Leica Microsystems) and assigned to four colonization classes: class I, no hyphal colonization; class II, ≤50% leaf area colonized by pathogen hyphae, without formation of conidiophores; class III, ≤75% leaf area colonized by hyphae, presence of conidiophores; class IV, >75% leaf area colonized by the pathogen, abundant conidiophores and sexual oospores. At least 150 leaves per were analyzed per genotype (Supplemental File 1). Statistically significant differences in *Hpa* colonization in each mutant were assessed with Chi-squared test in pairwise comparison with Wt (*p <*0.05).

For *Pst* infection, plants were spray-inoculated with bioluminescent *Pst::LUX* (Fan et al., 2008) at OD_600_ = 0.15 in 10mM MgSO_4_ + 0.01% v/v Silwet L-77. Inoculated plants were kept for three days at 100% RH to promote disease progression. After three days, digital photos of infected seedlings were taken with ChemiDoc XRS + Imager (BioRad). Relative bioluminescence was quantified as described previously (Furci et al., 2023; Furci et al., 2021). The day before inoculation, *Pst::LUX* bacteria were cultured from 1mL of frozen stocks in a shaking incubator (BioShaker BR-43FL) O/N at 28°C in King’s medium B, containing 50μg/mL Rifampicin and Kanamycin. More than 20 infected plants were scored per genotype (Supplemental File 1). Statistically significant differences in *Pst* colonization in each mutant were assessed with Student’s T-test in pairwise comparison with Wt (*α* =0.05).

*Pc* was maintained from frozen stock in 1/2 strength PDA plates kept for at least 2 weeks at room temperature in the dark until sporulation. Spores were gently scraped from plates using sterile water, and spore density was adjusted to 10^6^ spores/ml using a hemocytometer (Improved Neubauer, Hawksley, UK). For each seedling, the first two adult leaves were inoculated with 3µl droplets of the spore suspension, and kept at 100% RH following drop-inoculation. Six days after inoculation, infected leaves were collected for trypan blue staining and scored under steromicroscope (KL200 LED, Leica Microsystems). Infected leaves were assigned to four different classes based on size of the necrotic lesion caused by the pathogen, as described previously (Pastor-Fernández et al., 2020): class I, no necrotic lesion; class II, small necrotic lesion surrounding the drop-inoculation site.; class III, large necrotic lesion extending beyond drop-inoculation site; class IV, completely necrotic leaves. More than 35 leaves were scored per genotype (Supplemental File 1). Statistically significant differences in *Pc* necrotic lesion distribution in each mutant were assessed with Chi-squared test in pairwise comparison with Wt (*p <*0.05).

### RNA extraction, Illumina and Nanopore DRS sequencing

Total RNA was extracted from each genotype 48h after treatment with *Hpa* or Mock (sterile water), using Maxwell 16 LEV Plant RNA kit (Promega, USA). For each genotype/treatment combination, 12 seedlings were harvested in three biological replicates (four individual seedlings per sample).

For Illumina short-read sequencing, mRNA was isolated using NEBNext Poly(A) mRNA Magnetic Isolation Module (NEB). Libraries were prepared using NEBNext Ultra II RNA Library Prep Kit for Illumina (NEB) according to the manufacturer’s instructions, using size selection to obtain 450+ bp inserts. Concentration-normalized libraries were sequenced using Nova Seq 6000 (Illumina) at OIST sequencing facilities (150bp, paired-end)

For ONT-DRS sequencing, total RNA from Col-0 and *hakai-1* (Mock and *Hpa*) three biological replicates (same samples used for Illumina RNA-seq) were pooled into a single sample, and used to generate sequencing libraries with the Direct RNA Sequencing Kit (SQK-RNA002, Oxford Nanopore Technologies, UK) according to the manufacturer’s instructions. ONT-DRS was performed using MinION devices (device: MIN-101B, flow cells: FLO-MIN106 R9 version; Oxford Nanopore Technologies, UK) at our laboratory. We performed two sequencing runs for each condition. The basecalling was performed with Guppy (v4.4.2, https://nanoporetech.com/) to fit DENA requirement. We basecalled at least 2.6 million reads with a minimum of quality control of 8 for each of the four conditions.

### Illumina RNA-seq differential gene expression analysis and differential transcript usage analysis

Demultiplexed reads were trimmed using Trimmomatic (v0.39) (Bolger et al., 2014), with options LEADING:10 TRAILING:5 SLIDINGWINDOW:4:20 MINLEN:33. For Differential gene expression (DGE) analysis, paired trimmed reads were aligned to TAIR10 reference genome using Hisat2 (Kim et al., 2019) (default options), and read counts obtained with HT-seq “count” function (Putri et al., 2022) using Araport11 transcriptome as reference. DGE analysis was performed with DESeq2 (Love et al., 2014) in R (v4.3.1). For differential transcript usage (DTU) analysis, transcript-specific read counts were obtained by mapping paired trimmed reads to the Araport11 reference transcriptome using Salmon (Patro et al., 2017) (v1.10, mapping-based mode, default options). DTU analysis was performed on transcript-based Salmon counts using a published workflow (Love et al., 2018) that combines DRIMseq (Nowicka and Robinson, 2016) and DEXseq (Anders et al., 2012). For each indicated gene list, significant gene ontology (GO) enrichment for biological processes (BP) terms was performed with PANTHER (v18.0)(Thomas et al., 2022) using Benjamini-Hochberg false discovery rate (*fdr*) <0.05.

### m6A methylation calling from ONT-DRS and quantification

m6A methylation calling was performed with machine learning tool DENA for each of the four conditions (Wt Mock, Wt *Hpa*, *hakai-1* Mock and *hakai-1 Hpa*) using Araport11 transcriptome as reference, by following DENA instructions (Qin *et al*., 2022). Re-squiggling of ONT-DRS raw signal was done with Tombo (https://github.com/nanoporetech/tombo) as indicated in DENA manual, to obtain a unique mapping between the signal section of each base of each read and the reference sequence. ONT-DRS reads were aligned to the reference transcriptome using minimap2 (Li, 2018) with option: -ax map-ont -L --secondary=no. Next, the DENA scripts LSTM_extract.py and LSTM_predict.py were applied (Qin *et al*., 2022). For m6A methylation predictions, we only considered RRACH positions supported by at least 50 reads and with m6A ratio ≥0.1, as indicated by DENA’s authors (Qin *et al*., 2022). The m6A percentages was calculated as the number of modified reads divided by the number of total reads. To increase robustness in differential m6A calling, only m6A sites supported by at least 50 reads in each of the four conditions and methylated in Wt Mock (percentage ≥ 0.1) were considered (Supplemental Figure 2B). Because of DENA’s false positive rate (Qin *et al*., 2022), m6A sites with ratio below 0.1 were considered unmethylated and their ratio were set at 0. To identify differentially methylated m6A sites, Fisher’s exact test (*p* <0.05) was used for each condition in pairwise comparison with Wt Mock sample.

## Data availability

Illumina short-read RNA-seq data (adaptor-trimmed .fastq files) and ONT-DRS raw data (.fast5 files) generated in this study have been deposited in EMB-EBI European Nucleotide Archive database under accession codes PRJEB67534.

## Author contributions

L. F., J.B and H. S. designed the research and experiments. L. F. performed pathogen resistance assays and library preparation for Illumina RNA-seq and ONT-DRS. L. F. performed Illumina RNA-seq data analysis. J. B. performed ONT-DRS data analysis. L. F., J. B. and H. S. wrote and edited the manuscript.

## Acknowledgments

We thank Prof. Rupert G. Fray and associated lab members for kindly providing *hakai-2* and *mta* mutant Arabidopsis seeds. We are grateful for the help and support provided by the Sequencing Section (SQC) of Research Support Division at OIST Graduate University for performing Illumina short-read mRNA sequencing. This work was supported by MEXT Grant-in-Aid for Transformative Research Areas (A) JP20H05913 to H. S. and by JSPS Grant-in-Aid for Early-Career Scientists 23K13957 to L. F.

## Supplemental Figures

**Supplemental Figure 1:**
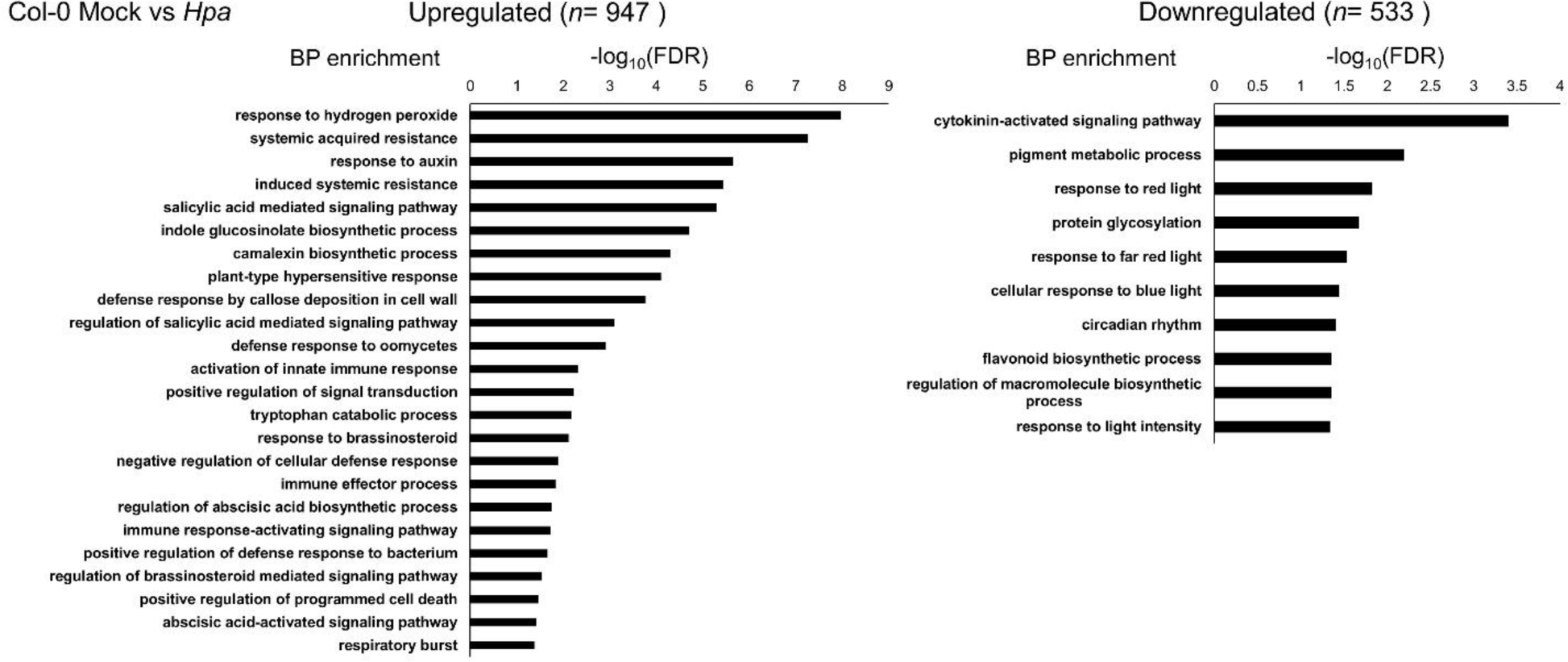
Gene ontology term enrichment (biological processes) for differentially upregulated and downregulated genes in *Hpa*-treated Wt seedlings compared to Mock-treated ones.

**Supplemental Figure 2:**
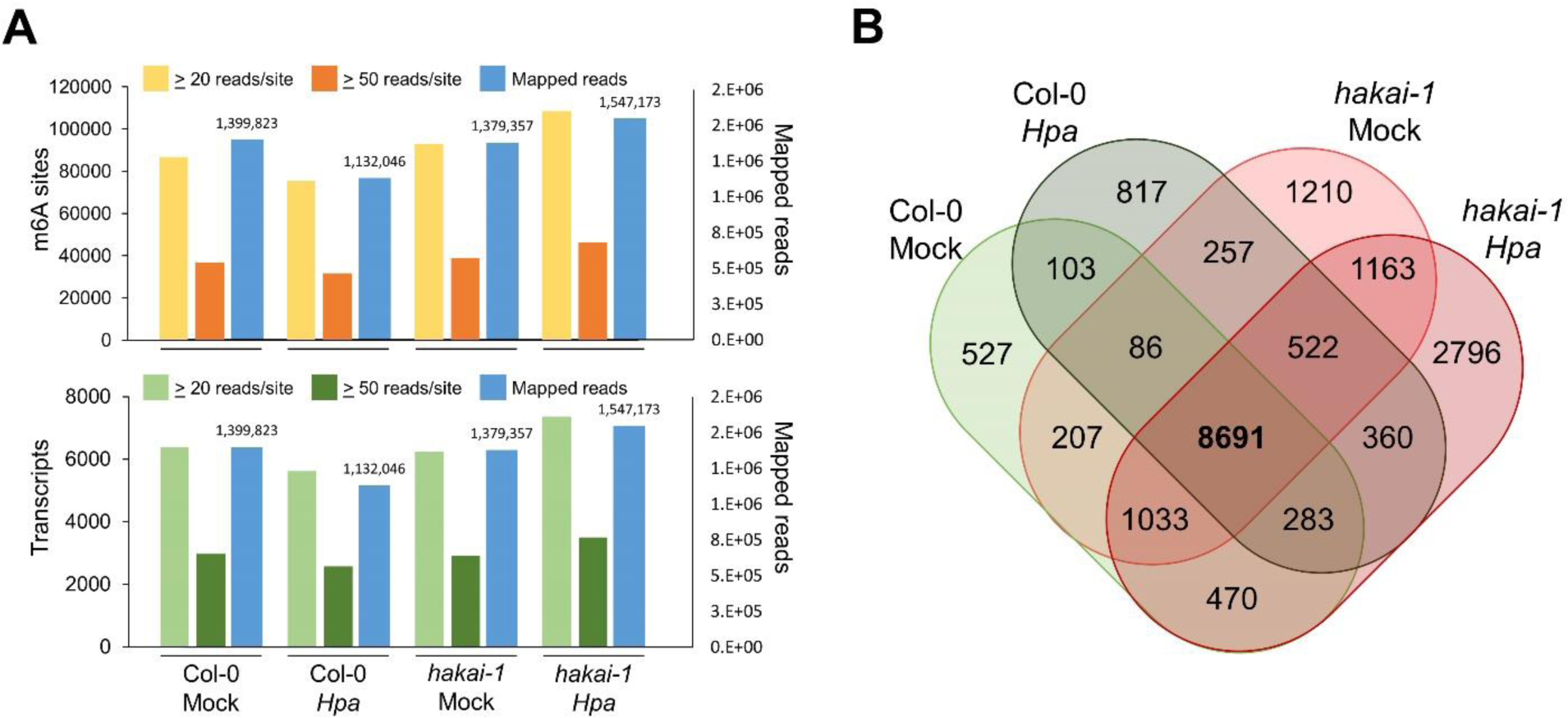
**A:** correlation between number of identified m6A sites (top panel, yellow/orange bars) or number of identified transcripts covered by at least one m6A site (bottom panel, light/dark green bars) at different coverage thresholds with number of mapped reads (blue bars) in each condition. **B:** overlap of m6A sites (50 reads coverage threshold) identified in each condition. Center overlap in bold indicates a subset of 8691 sites covered by at least 50 reads in all four conditions.

**Supplemental Figure 3:**
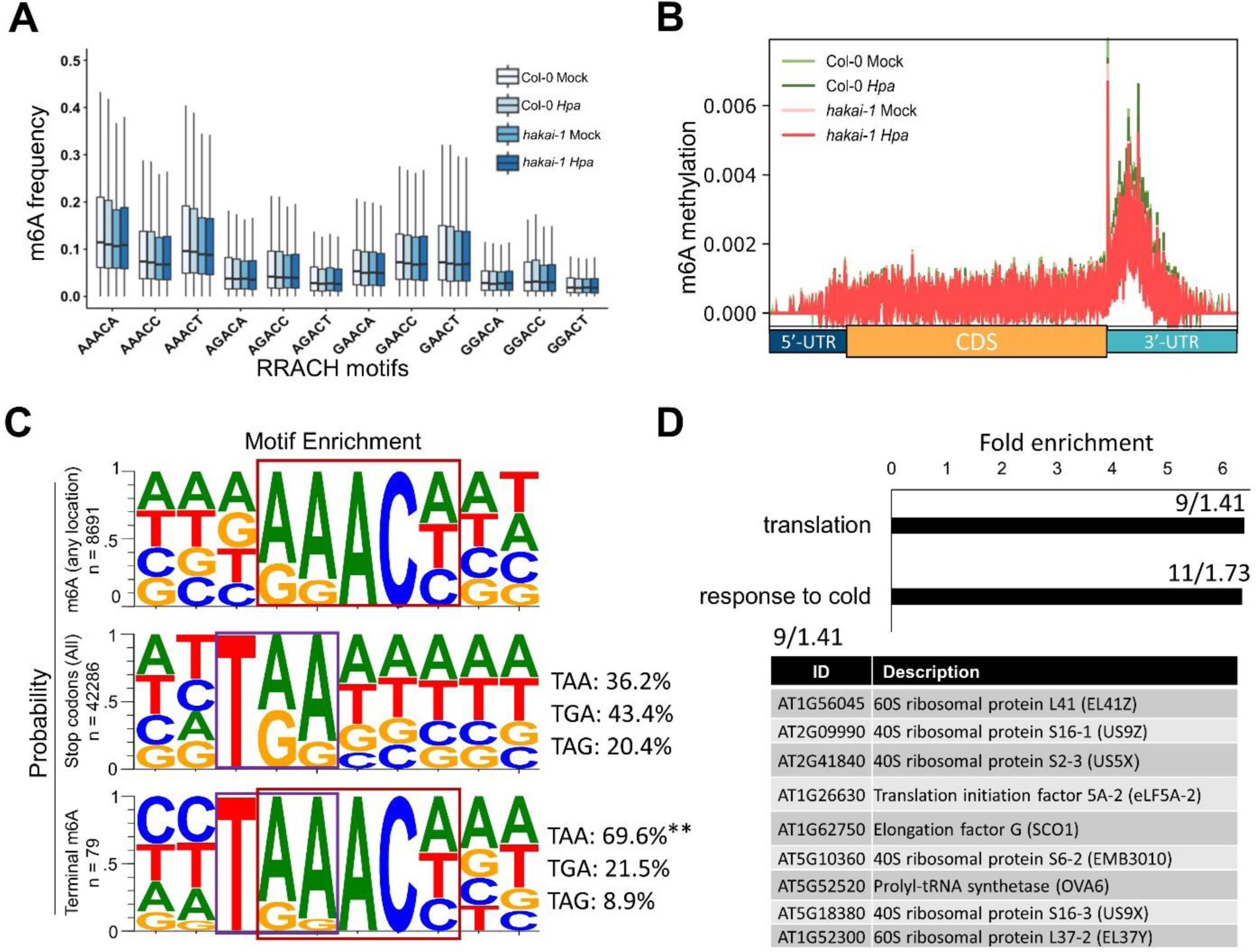
**A:** m6A methylation frequency for each of 12 possible RRACH motifs across conditions**. B:** Mean m6A methylation ratio at 1bp resolution over transcript regions for shared m6A sites (*n*= 8691) across conditions. **C:** motif enrichments at 10 nucleotide region for shared m6A sites (top), stop-codon region of all Araport11 transcripts (mid) and 79 stop-codon terminal m6A sites (bottom). Red boxes indicate RRACH motif and purple boxes indicate stop-codon. Statistically significant differences in stop-codon nucleotide triplet frequencies between all transcripts (mid) and stop-codon terminal m6A sites (bottom) were assessed by Chi-squared test (**, *p* < 0.01). **D:** Top panel: GO terms enrichment (BP) for the 73 transcripts with stop-codon terminal m6A. For each enriched term, the number of genes identified/expected number is indicated above. Bottom panel: list and description of the nine genes with stop-codon terminal m6A associated with the “translation” GO term.

**Supplemental Figure 4:**
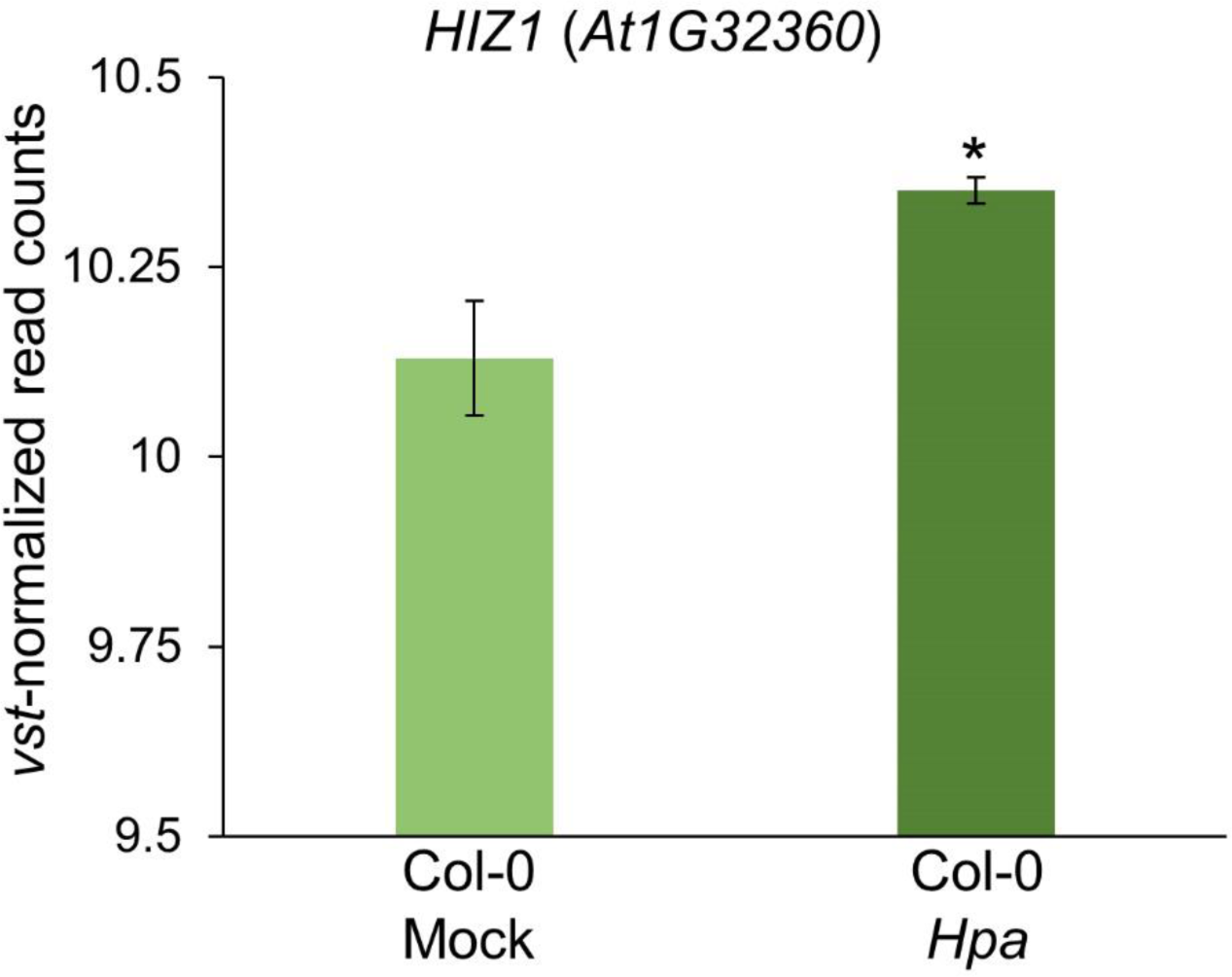
Average *vst*-normalized read counts(Love *et al*., 2014) from Illumina RNA-seq for *HAKAI-INTERACTING ZINC FINGER PROTEIN 1* (*HIZ1*, *At1G32360*) between Mock- and *Hpa-*treated Col-0 seedling. Error bars represent standard error of the mean for three biological replicates. Statistically significant differences in read counts were calculated with “*DESeq2*” R package (Wald test, *q*= 0.014)

**Supplemental Figure 5:**
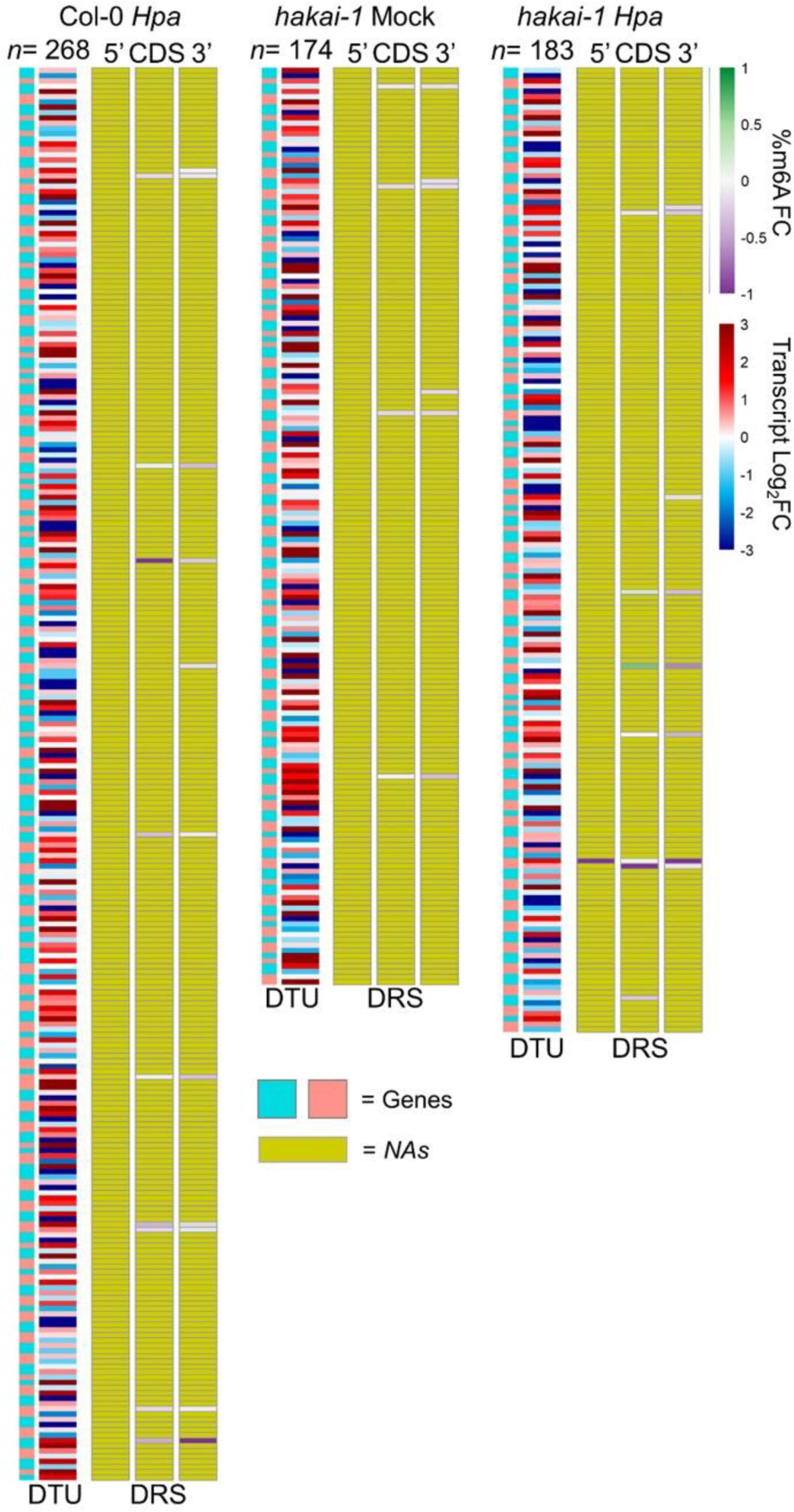
comprehensive overview of all genes showing differential transcript usage (DTU) in each condition (compared to Mock-treated Col-0). DTU heatmap displays log_2_ fold changes in isoform expression for each identified gene (cyan and pink annotation); DRS heatmap shows fold changes in m6A percentage across transcript region of each isoform (NA = no data available).

